# On the Optimal Use of Isotherm Models for the Characterization of Biosorption of Lead Onto Algae

**DOI:** 10.1101/017426

**Authors:** F. Brouers, Tariq J. Al-Musawi

## Abstract

In this study the experimental isotherm data of biosorption of Pb(II) onto algae was modeled using several models. These models are: Langmuir, Hill-Sips, Brouers-Sotolongo, Brouers-Gaspard, and Redlich-Peterson models. The coefficients of each model were determined by non-linear fitting using Mathematica9 program. The maximum Pb(II) removal rate increased with the increase of temperature and reached the maximum value (98%) at the temperature of 40 °C. Even if the R^2^ error quantity is not the unique and always the best measure for nonlinear fitting, the Brouers-Sotolongo model gives in all cases the best fit and is definitely the most suitable one to satisfactorily describe bioisorption of Pb(II) on the algal biomass. In addition, this study shows that a complete set of data is necessary to have a good representation of the isotherm.

## 1. Introduction

Biosorption is a process which utilizes inexpensive dead biomass to extract toxic heavy metals from aqueous solutions (Kratochvil and Volesky, 1998). Biosorption is proven to be quite effective at removing metals ions from contaminated solutions in a low cost and environment-friendly manner. Various materials such as algae (Sulaymon et al., 2013^a^), fungi (Holan and Volesky, 1995), grass (Sulaymon et al., 2014) and agriculture waste (Hameed et al., 2008; Rathinam et al., 2010) have been used to sorb metal ions from the aqueous solution. The term algae refers to a large and diverse assemblage of organisms that contain chlorophyll and carry out oxygenic photosynthesis (Davis et al., 2003). Algae are one of the most abundant and highly available natural resources in tropical ecosystems. Cyanophyta (blue–green algae), Chlorophyta (green algae), Rhodophyta (red algae), and Phaeophyta (brown algae) are the divisions of the largest visible algae. Many studies concluded that the algal biomass is a very promising material to be used as biosorbent to remove various kinds of pollutants from contaminated water and wastewaters (Sulaymon et al., 2013^b^; Romera et al., 2007; Altenor, et al., 2012). This can be attributed that the algal biomass has many negative charge active groups on its surface cell wall such as hydroxyl, carboxyl, amino, sulfhydryl, and sulfonate (Davis et al., 2003).

The term isotherm is used for describing the retention of a substance on a solid at a constant temperature. The isotherm study is a major tool to predict the efficiency of a sorbent to remove a given pollutant from polluted water (Ncibi et al., 2008). Analysis of experimental isotherm data by adapting to different isotherm models is an important step to develop an equation which accurately represents the results and which can be used for design purposes. Hence, a large number of studies have been done to develop a mathematical isotherm models and to verify their suitability for describing biosorption data of heavy metals by algal biomass data as well as to understand the sorption isotherm phenomenon between the biomass surface and the metal molecules. Most of these models are empirical and bring little information on the physicochemical processes responsible for the particular shape of the isotherm curves. Brouers (2014 ^a^) has shown that some of the most empirical models that can be used to describe the isotherm data were approximations of a generalized Brouers-Sotolongo model. This model is derived from the Burr-Madalla distribution which is itself solution of a birth and death differential equation which can describe sorption-desorption mechanisms (Burr, 1942; Maddala, 1996). The author concluded that only the Langmuir, the Hill-Sips and the Brouers-Sotolongo isotherms were genuine statistical functions and had the correct asymptotic limits for low and high concentrations. In addition, the author pointed out that a statistical analysis had a meaning only if the experimental data were complete until the saturation of the sorption. Furthermore, Ion-exchange is an important concept in biosorption mechanisms using algae, because it explains many of the observations made during heavy metal uptake experiments. In this case, the ion exchange reaction type occurs between light metals already bound to the algae and other metals present in the aqueous solution (Naja and Volesky, 2006). The ion-exchange model is certainly a better representation of the biosorption process using algae since it reflects the fact that most algal biomass is either protonated or contains light metal ions such as K^+^, Na^+^ and Mg^2+^, which are released upon binding of a heavy metal cation. It should be pointed out that the ion exchange model does not explicitly identify the binding mechanism (Davis et al., 2003). Hence, verification of the various isotherm models is a growing area of study.

Despite the fact that linear regression is still frequently used (Sulaymon et al., 2010), non-linear analysis of isotherm data is an interesting and useful mathematical approach for describing biosorption isotherms for water and wastewater treatment applications and to predict the overall sorption behaviour under different operating conditions (Ho, 2006). Indeed, the linearization of non linear isotherm equations has several disadvantages. This process implicitly alters their error structure and may also violate the error variance and normality assumptions of standard least squares (Ratkowsky, 1990). In general, non linear regression gives a more accurate determination of model parameters than linear regression method. In recent years, several error analysis methods, such as the coefficient of determination (R^2^), the sum of the errors squared, a hybrid error function, Marquardt’s percent standard deviation, the average relative error and the sum of absolute errors, have been used to determine the best-fitting isotherm (Ho et al., 2002; Allen et al., 2003). Ho (2004) point out that it is not appropriate to use the R^2^ of linear regression analysis for comparing best fitting of models since this parameter failed to represent the error for fitting with linear equation. As different forms of the equations affected R^2^ values more significantly during the linear analysis, the non-linear analysis might be a method of avoiding such errors (Ho and Wang, 2004; Kumar and Sivanesan, 2005). Additionally, Brouers (2014^b^) showed that the use of nonlinear programs allows an easy and precise fitting of the experimental data with the theoretical models.

The aim of this paper is to apply and demonstrate the interest of new methodology introduced by Brouers (2013) and Brouers (2014^a^) on the isotherm data for the biosorption of Pb(II) in aqueous solution using algae. The non linear method of five isotherm model, Langmuir, Hill-Sips, Brouers-Sotolongo, Brouers-Gaspard, and Redlich-Peterson models, were compared with the experimental data. A trial-and-error procedure was used for the non-linear analysis method using the Mathematica 9 program and based on the Burr-Madalla distribution function introduced by Burr (1942).

## 2. Materials and Methods

### 2.1 Materials

Fresh mixture of green and blue-green algae was used in this study as a biosorbent. This material was collected from the artificial irrigation canal in Baghdad University, Iraq. It was mainly a mixture of three species of algae. Blue-green Oscillatoria princeps alga was the highest percentage (88%), green Spirogyra aequinoctialis alga was (9%), and blue-green Oscillatoria subbrevis alga (3%). And to make it user friendly the collected algae were not separated. The foreign matters were removed manually from the collected algae, then rinsed with tap and distilled water to remove of dirt, sands, and external salts. Afterward, the washed algae were kept in air for removing water and dried at an oven temperature of 65 °C for 48 h. The dried biomass were roughly chopped, grounded into powder, sieved, and kept in air-tight polyethylene container at room temperature. An average size of 0.54 mm was used for biosorption experiments with required amounts.

All the chemicals used in this work are analytical grade reagents with deionized water used for solutions preparation. Stock solution (1000 mg/l) of Pb(II) was prepared by dissolving the appropriate weight of lead chloride in distilled water and kept in glass container at room temperature. The desired concentrations were prepared by diluting the stock solution in accurate proportions to different initial concentrations. The initial pH of the working solutions was adjusted by addition of 1 mol/l NaOH or HCl using a pH meter (WTW, inoLab 720, Germany). All the glassware used for dilution, storage and experimentation were cleaned with detergent, thoroughly rinsed with tap water, soaked overnight in a 20% HNO_3_ solution and finally rinsed with distilled water before use. Table (1) listed the properties of algal biomass and Pb(II) salt used in this study.

**Table (1):**
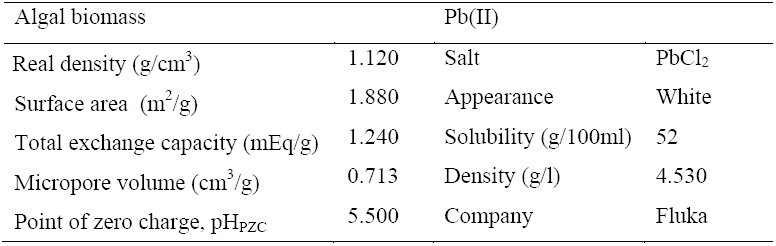
Properties of the algal biomass and Pb(II) salt used in this study.

| Algal biomass |  | Pb(II) |  |
| --- | --- | --- | --- |
| Real density (g/cm<sup>3</sup>) | 1.120 | Salt | PbCl<sub>2</sub> |
| Surface area (m<sup>2</sup>/g) | 1.880 | Appearance | White |
| Total exchange capacity (mEq/g) | 1.240 | Solubility (g/100ml) | 52 |
| Micropore volume (cm<sup>3</sup>/g) | 0.713 | Density (g/l) | 4.530 |
| Point of zero charge, pH<sub>PZC</sub> | 5.500 | Company | Fluka |

### 2.2 Methods

#### 2.2.1 Experiments

Isotherm experiments were carried out in 250 ml stoppered conical flasks containing 0.05, 0.1, 0.3, 0.5, 0.8, 1, 2, and 3 g of algal biomass and 100 ml of Pb(II) solution. These experiments were performed at the same initial concentration (50 mg/l) for Pb(II) and repeated for temperature range from 10 to 40 °C. According to the previous studies, the optimum pH for Pb(II) removal using algae is in the range between pH 2 and pH 4 (Sulaymon et al., 2013^a^; Davis et al., 2003). In addition, several authors showed that further increases in temperature (above 40 °C) lead to a decrease the percentage removal. This may be attributed to an increase in the relative desorption of the metal from the solid phase to the liquid phase, deactivation of the biosorbent surface, destruction of active sites on the biosorbent surface **(Saleem, et al., 2007; Meena, et al., 2005.** Hence, in this study all the isotherm experiments were conducted at temperature below 40 °C and the pH solution was adjusted to the 3. The flasks were placed in a shaker (Edmund Buhler, 7400 Tubingen Shaker-SM 25) with constant shaking at 200 rpm for 4 h. After equilibrium condition, the sorbent was separated from aqueous solution by using filter paper (WHATMAN, No.42, diameter 7 cm). The Pb(II) concentrations in both initial and withdrawn samples were determined using atomic absorption spectrophotometer (type: SHIMADZU, AAS 7200, Japan). Each sample was measured thrice, and the results were given as the average value. Biosorption capacity at equilibrium conditions and the percentage of removal (%) were calculated using eq(1) and eq(2), respectively.

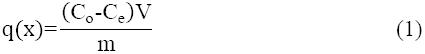

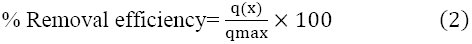

where, q(x) and are the equilibrium and maximum biosorption capacity of Pb(II) (mg/g), respectively; C_o_ and C_e_ are the initial and equilibrium Pb(II) concentrations in the aqueous solution (mg/l), respectively; V is the volume of used solution (l); and m is the mass of used adsorbent (g).

#### 2.2.2 Isotherm models

There are several isotherm models with two or more than two parameters available for analyzing the experimental parameters. The parameters of each model often provide insights into the sorption mechanism, the surface properties and the affinity of the sorbent (Yu and Neretnieks, 1990). In the present study, five different isotherm models were tested under different adsorption temperatures. These models are: Langmuir, Sips, Brouers-Sotolongo, Brouers-Gaspard, and Redlich-Peterson models were chosen to fit the experimental data and were derived from the General Brouers-Sotolongo equation as will be shown below (Brouers, 2014^a^). The non linear regression analysis using Mathematica program, version 9, was used for direct determination of each model parameters. In this case, a trial-and-error procedure, which is applicable to computer operation, was developed to determine the isotherm parameters using an optimization routine to maximize the coefficient of determination (R^2^) between the experimental data and isotherms. The applicability and suitability of the isotherm equation to the equilibrium data were compared by judging the values of the R^2^ values.

The General Brouers-Sotolongo (GBS) isotherm is represented by the following equation (Brouers, 2014^a^):

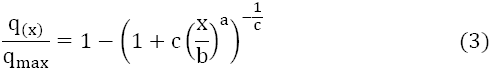

Where q_max_ is the maximum sorbed quantity (mg/g), x is the residual metal concentration in solution at equilibrium (mg/l), b is the isotherm constant (mg/l), a is a coefficient depends on the fractal nature of the system and this exponent is a measure of the width of the adsorption energy distribution and energy heterogeneity of the sorbent surface (Ahmad et al., 2014), and c is the coefficient related to its cluster organization (Stanislavsky and Weron, 2013).

If c=1 and a=1, GBS reduces to the Langmuir model (eq. (4)). Langmuir model is the best known and most often used isotherm model for the sorption of a solute from a liquid solution. This model is based on the assumption that sorption sites are identical and energetically equivalent, and that the solute is immobilized under the form of monolayer coverage (**Langmuir, 1918**).

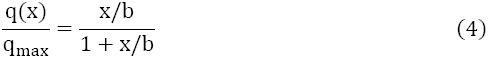

**Hill-Sips** (HS) isotherm model (eq.(5)) can be derived from GBS when c=1. Hill-Sips isotherm model is a combined form of Langmuir and Freundlich expressions deduced for predicting the heterogeneous adsorption systems and circumvent the limitation of the rising adsorbate concentration associated with Freundlich isotherm model. At high adsorbate concentration, it predicts monolayer adsorption characteristics of Langmuir, while in low adsorbate concentration, it reduces to Freundlich isotherm (Sips, 1948).

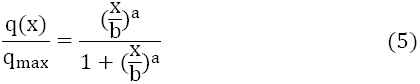

For c=0, one recovers the Brouers-Sotolongo (BS) isotherm equation (eq. (6)). Brouers-Sotolongo (BS) is one of the earlier attempts to formulate a theory of sorption on a non-uniform surface (Ncibi, et al., 2008). It has been proposed to analyse sorption processes on highly heterogeneous systems. The surface of the biosorbent is assumed to be made of a finite number of patches of sites of different sorption energies (Brouers et al., 2005).

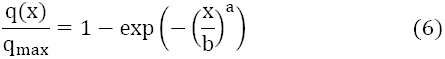

Indeed, it is difficult in many cases to choose between HS and BS isotherms, it is why it has been proposed to use in these cases an intermediate value for c (c=1/2) and introduce a new intermediate formula which is named Brouers-Gaspard (BG) isotherm equation (eq.(7)).

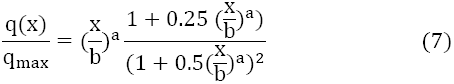

All these equations have good physical asymptotic behaviors:

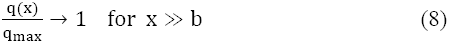

and

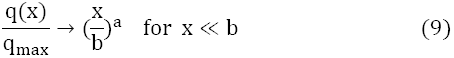

Another commonly used empirical isotherm is the Redlich-Peterson isotherm (RP) (eq. (10)). Redlich-Peterson isotherm model incorporates the features of the Langmuir and Freundlich models into a single equation and presents a general isotherm. This model can be applied to express the sorption process when dealing with a certain pollutants at high concentration (Quintelas, et al., 2008).

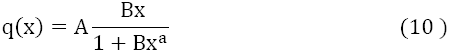

This equation does not have the physical correct asymptotic limit since it gives:

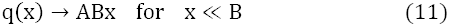

and

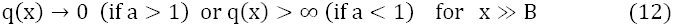

Where A and B are the Redlich–Peterson isotherm constants.

## 3. Results and discussion

The bisorption isotherm of Pb(II) onto algal biomass was investigated as a function of temperature and the results are depicted in Fig. 1 (a). The graph are plotted in the form of Pb(II) removal efficiency against algal biomass weight. These data may help in understanding the controlling mechanisms and quantifying the sorption properties of the sorbent, maximum sorption capacity and affinity of the sorbent for the target metal (Langmuir, 1918). For all experimental data shown in this figure, it can be seen that the removal efficiency was increased with the increase of algal biomass weight from 0.05 to 1 g, implying that the optimum amount of biosorbent is 1 g algal biomass/200 ml solution, then beyond 1 g algal biomass dose the removal efficiency reached a plateau which demonstrating the equilibrium state. The reason being that an increase in the biosorbent quantity in the aqueous solution results in a larger exchangeable sites or surface area for Pb(II) sorption, hence the rise in the removal percentage (Hekmatzadeh, et al., 2013). By further increment in sorbent dose, the removal capacity was not increased possibly due to the aggregation of available binding sites. Moreover, Fig. 1 (a) shows that with an increase in temperature, the percentage of Pb(II) removal increases and the maximum removal efficiencies were obtained at 40 °C. This means that Pb(II) binding on active sites of the biosorbent becomes stronger at higher temperature and that the sorption process is endothermic. This can be attribute to the increase in temperature is knowing to increase the rate of diffusion of the adsorbate molecules across the external boundary layer and in internal pores of the adsorbent particles as a result of the reduced viscosity of the solution (Bulut et al., 2012). A similar trend was obtained Sulaymon et al., (2013^b^).

**Fig. 1.**
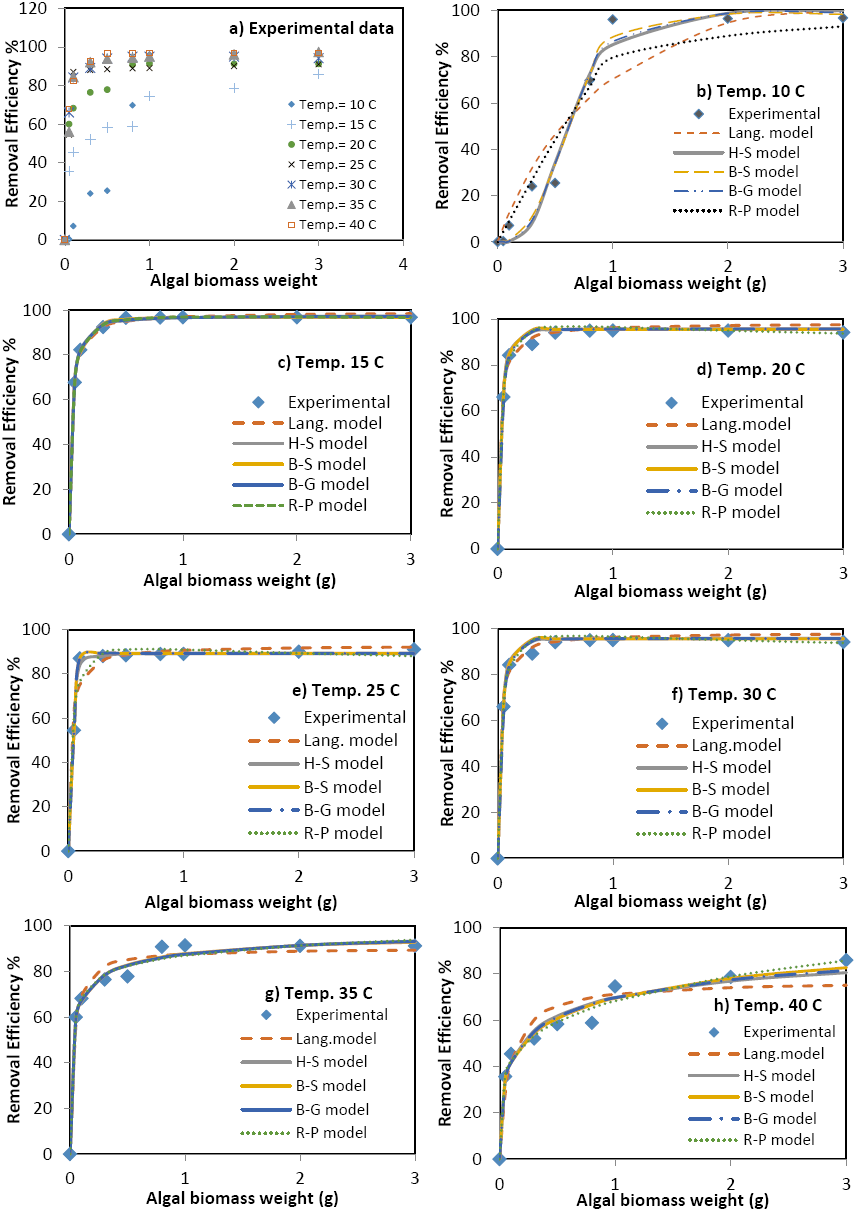
Experimental data of biosorption isotherm of Pb(II) ions at different temperatures using algal biomass and simulation of isotherm data with several models

The experimental isotherm data have been systematically modeled by the aforementioned isotherm models and the results are depicted in Fig. 1(b-h). The parameters of these models and the coefficient of determination (R^2^) values are listed in Tables 2. Based on the analysis of the estimated variance (R^2^ values), the B-S model gave a best fit of experimental data for all temperature values. The R^2^ values of Brouers–Sotolongo model were much closer to 1.0 than the values obtained with the other models. Such fitting tendency would refer to the presence of active sites with heterogeneous sorption interactions (Ncibi, et al., 2008; Altenor et al., 2012). In order to find out why this newly established Brouers–Sotolongo isotherm model was able to fit better the equilibrium data than the widely used models, a couple of points will be detailed hereafter. First of all, the B-S isotherm was an initiative to propose a new equation able to describe the adsorption process considering from the start that it is a complex system. Such trend would probably enlighten the energetic heterogeneity at the interface algal biomass. The heterogeneous surface stems from two sources known as geometrical and chemical ones. Geometrical heterogeneity is a result of differences in size and shape of pores, cracks and pits. Chemical heterogeneity is associated with different functional groups such as hydroxyl, amine, and carboxyl and aldehydes determining the apparent chemical character of an algal biomass surface (Sulaymon et al., 2013^a^; Kratochvil and Volesky, 1998). Indeed, under the used experimental conditions, the interactions involved in such bisorption case would be between lead molecules and algal biomass cell surfaces via different possible interactions mainly electrostatic attraction, ion exchange and complexation between lead and hydroxyl, amine, and carboxyl groups (Davis et al., 2003). Thus, such tendency would lead to a highly heterogeneous sorption energy landscape, explaining therefore the good fit of the B-S isotherm model.

**Table (2).**
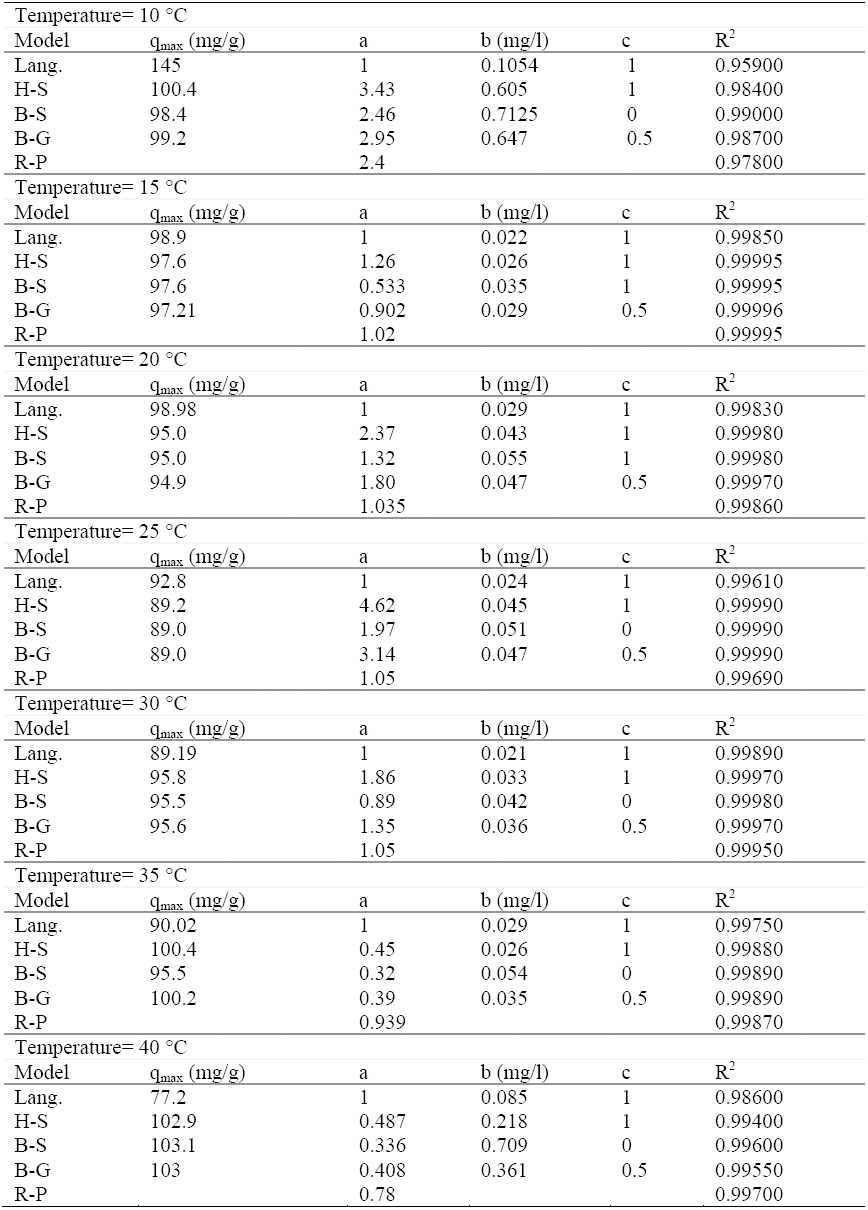
Isotherm modeling results related to the biosorption of Pb(II) onto algal biomass

| Temperature= 10 °C |  |  |  |  |  |
| --- | --- | --- | --- | --- | --- |
| Model | $q_{\max}$ (mg/g) | a | b (mg/l) | c | $R^2$ |
| Lang. | 145 | 1 | 0.1054 | 1 | 0.95900 |
| H-S | 100.4 | 3.43 | 0.605 | 1 | 0.98400 |
| B-S | 98.4 | 2.46 | 0.7125 | 0 | 0.99000 |
| B-G | 99.2 | 2.95 | 0.647 | 0.5 | 0.98700 |
| R-P |  | 2.4 |  |  | 0.97800 |
| Temperature= 15 °C |  |  |  |  |  |
| Model | $q_{\max}$ (mg/g) | a | b (mg/l) | c | $R^2$ |
| Lang. | 98.9 | 1 | 0.022 | 1 | 0.99850 |
| H-S | 97.6 | 1.26 | 0.026 | 1 | 0.99995 |
| B-S | 97.6 | 0.533 | 0.035 | 1 | 0.99995 |
| B-G | 97.21 | 0.902 | 0.029 | 0.5 | 0.99996 |
| R-P |  | 1.02 |  |  | 0.99995 |
| Temperature= 20 °C |  |  |  |  |  |
| Model | $q_{\max}$ (mg/g) | a | b (mg/l) | c | $R^2$ |
| Lang. | 98.98 | 1 | 0.029 | 1 | 0.99830 |
| H-S | 95.0 | 2.37 | 0.043 | 1 | 0.99980 |
| B-S | 95.0 | 1.32 | 0.055 | 1 | 0.99980 |
| B-G | 94.9 | 1.80 | 0.047 | 0.5 | 0.99970 |
| R-P |  | 1.035 |  |  | 0.99860 |
| Temperature= 25 °C |  |  |  |  |  |
| Model | $q_{\max}$ (mg/g) | a | b (mg/l) | c | $R^2$ |
| Lang. | 92.8 | 1 | 0.024 | 1 | 0.99610 |
| H-S | 89.2 | 4.62 | 0.045 | 1 | 0.99990 |
| B-S | 89.0 | 1.97 | 0.051 | 0 | 0.99990 |
| B-G | 89.0 | 3.14 | 0.047 | 0.5 | 0.99990 |
| R-P |  | 1.05 |  |  | 0.99690 |
| Temperature= 30 °C |  |  |  |  |  |
| Model | $q_{\max}$ (mg/g) | a | b (mg/l) | c | $R^2$ |
| Lang. | 89.19 | 1 | 0.021 | 1 | 0.99890 |
| H-S | 95.8 | 1.86 | 0.033 | 1 | 0.99970 |
| B-S | 95.5 | 0.89 | 0.042 | 0 | 0.99980 |
| B-G | 95.6 | 1.35 | 0.036 | 0.5 | 0.99970 |
| R-P |  | 1.05 |  |  | 0.99950 |
| Temperature= 35 °C |  |  |  |  |  |
| Model | $q_{\max}$ (mg/g) | a | b (mg/l) | c | $R^2$ |
| Lang. | 90.02 | 1 | 0.029 | 1 | 0.99750 |
| H-S | 100.4 | 0.45 | 0.026 | 1 | 0.99880 |
| B-S | 95.5 | 0.32 | 0.054 | 0 | 0.99890 |
| B-G | 100.2 | 0.39 | 0.035 | 0.5 | 0.99890 |
| R-P |  | 0.939 |  |  | 0.99870 |
| Temperature= 40 °C |  |  |  |  |  |
| Model | $q_{\max}$ (mg/g) | a | b (mg/l) | c | $R^2$ |
| Lang. | 77.2 | 1 | 0.085 | 1 | 0.98600 |
| H-S | 102.9 | 0.487 | 0.218 | 1 | 0.99400 |
| B-S | 103.1 | 0.336 | 0.709 | 0 | 0.99600 |
| B-G | 103 | 0.408 | 0.361 | 0.5 | 0.99550 |
| R-P |  | 0.78 |  |  | 0.99700 |

It is worth to note that using only R^2^ to determine the best-fitting model is not sufficient and could lead to some ambiguities when the set of data is not complete. Indeed, the results in Table 2 showed that based on R^2^ values, H-S, B-G and R-P models seem to be adequate and the parameters of these models were the best fitting for the experiments results. It must be noted however that the R-P model which does not have the proper asymptotic behaviors is generally less performing than the other isotherm here that we have complete data. The calculation of the error deviation using another methods such as the average relative error and/or the Marquardt’s percent standard error may be also of interest to describe the validity of sorption model as suggested by Ncibi et al. (2008) for the adsorption of phenol and methylene blue (MB), respectively onto a non-porous adsorbent, Posidonia oceanica fibers and two porous adsorbents, chemically and physically activated carbons prepared from vetiver roots.

## 4. Conclusions

This study revealed the following conclusions:

- This study confirmed that algal biomass is a promising biosorbent for Pb(II) removal from aqueous solution and the maximum removal efficiency was reach 98% at 40 °C and pH 3.
- Excellent results were obtained by using original non linear isotherm models and one should leave the old-fashion linear methods now that one has rapid computers and that performing nonlinear recursion methods are on the market.
- The experimental isotherm data were better described by the Brouers-Sotolongo model involving the biosorption of Pb(II) on a heterogeneous surface onto algal biomass particles.
- This study shows that a complete set of data is necessary to have a good representation of the isotherm.
- Using only coefficient of determination is not always an appropriate tool to compare the goodness of the non linear fit of an isotherm models.

## Acknowledgments

We would like to express our sincere thanks and deep gratitude to Professor Sarra Gaspard (France) and Professor Mongi Seffen (Tunis) for their useful discussions on the use of the non-linear approach. Special thanks to the Environmental Engineering Department, University of Baghdad, Iraq for supporting this work.

## References

1 Ahmad M.Z., Ahmad N., Bello O.S., Modified durian seed as adsorbent for the removal of methyl red dye from aqueous solutions, Appl Water Sci, 2014, DOI 10.1007/s13201-014-0208-4.

2 Allen SJ, Gan Q, Matthews R, Johnson PA. Comparison of optimised isotherm models for basic dye adsorption by kudzu. Bioresour Technol 2003;88(2):143–52.

3 Altenor S., Ncibi M.C., Emmanuel E., Gaspard S., Textural characteristics, physiochemical properties and adsorption efficiencies of Caribbean alga Turbinaria turbinata and its derived carbonaceous materials for water treatment application, Biochemical Engineering J. 67(2012) 35–44.

4 Brouers, F. (2013), Sorption Isotherms and Probability Theory of Complex Systems. arXiv: 1309, 5340–55.

5 Brouers, F., Sotolongo O., Marquez F.,Pirard J. P., Microporous and heterogeneous surface adsorption isotherms arising from Levy distributions, Phys. A 349 (2005) 271.

6 Brouers, F. (2014a), Statistical Foundation of Empirical Isotherms. Open Journal of Statistics, 4(9), 687.

7 Brouers, F. (2014b), The Fractal (BSf) Kinetics Equation and Its Approximations. Journal of Modern Physics, 5(16), 1594.

8 Bulut Y., Gul A., Baysal Z., Alkan H., 2012. Asorption of Ni(II) from aqueous solution by Bacillus subtilis, Desalination and Water Treatment 49, 74–80.

9 Burr, I.W. (1942) Cumulative Frequency Functions. The Annals of Mathematical Statistics, 13, 215–232.

10 Davis A., Volesky B., Mucci A., A review of the biochemistry of heavy metals biosorption by brown algae, J. Water Res. 37 (2003) 4311–4330.

11 Hekmatzadeh A., Karimi-Jashni A., Talebbeydokhti N., Klove B., Adsorption Kinetics of Nitrate Ions on Exchange Resin, Desalination 326(2013) 125–134.

12 Hameed, B.H., Mahmoud D.K., and Ahmed, A.L., 2008, Sorption Equilibrium and Kinetics of Basic Dye from Aqueous Solution Using Banana Stalk Waste, J. Hazar. Mater., Vol. (158), PP. 499–506.

13 Ho Y-S, Selection of optimum sorption isotherm, Carbon 42 (2004) 2115–2116.

14 Ho Y-S, Wang C.C, Pseudo-isotherms for the sorption of cadmium ion onto tree fern, Process Biochem. 39 (2004) 759.

15 Ho Y-S, Porter JF, McKay G. Equilibrium isotherm studies for the sorption of divalent metal ions onto peat: copper, nickel and lead single component systems. Water Air Soil Pollut 2002; 141(1–4): 1–33.

16 Ho Y-S., Isotherms for the Sorption of Lead onto Peat: Comparison of Linear and Non-Linear Methods, Polish Journal of Environmental Studies 15 (2006) 81–86.

17 Holan ZR, Volesky B (1995) Accumulation of cadmium, lead and nickel by fungal and wood biosorbents. J Appl Biochem Biotechnol 53:133–142.

18 Kratochvil D., Volesky B., Advances in biosorption of heavy metals, J. Trend Biotechnol. 16 (1998) 291–300.

19 Kumar K.V., Sivanesan S., Prediction of optimum sorption isotherm: comparison of linear and non-linear method, J. Hazard. Mater. B126 (2005) 198.

20 Langmuir I., (1918) The adsorption of gases on plane surfaces of glass, mica and platinum. J Am Chem Soc 40:1361–1402.

21 Maddala, G.S. (1996) Limited -Dependent and Qualitative Variables in Econometrics Cambridge University Press.

22 Meena, A.K, Mishra, G.K., Rai, P.K., Rajagopal. C., and Nagar, P.N., 2005, Removal of Heavy Metal Ions From Aqueous Solutions Using Carbon Aerogel as an Adsorbent, J. Hazard. Mat., Vol. (122), PP: 161–170.

23 Naja G, Volesky B (2006) Behavior of the mass transfer zone in biosorption column. J Environ Sci Technol 40:3996–4003

24 Ncibi M.C., Altenor S., Seffen M., Brouers F., Gaspard S., Modelling single compound adsorption onto porous and non-porous sorbents using a deformed Weibull exponential isotherm, Chem. Eng. J. 145 (2008) 196–202.

25 Ncibi, M.C., Majoub, B., Steffen, M., F. Brouers, F. and Gaspard, S. (2009) Sorption Dynamic Investigation of Chromium (VI) onto Posidonia Oceanica Fibres: Kinetic Modelling Using New Generalized Fractal Equation. Biochemical Engineering Journal, 46, 141–46.

26 Quintelas, C., Fernandes, B., Castro, J., Figueiredo, H., and Tavares, T. 2008, Biosorption of Cr(VI) by three different bacterial species supported on granular activated carbon—a comparative study, J. Hazard.Mater., Vol. (153), PP. 799–809.

27 Rathinam, B. Maharshi, S.K. Janardhanan, R. Jonnalagadda, B.U. Nair, Biosorption of cadmium metal ion from simulated wastewater using hypnea Valentiae biomass: A kinetic and thermodynamic study, J. Bioresour. Technol. 101 (2010) 1466–1470.

28 Ratkowsky DA. Handbook of nonlinear regression models. New York: Marcel Dekker Inc; 1990.

29 Redlich, O. and Peterson, D. L. (1959). A Useful Adsorption Isotherm. Journal of Physical Chemistry, 63, 1024–1024.

30 Romera E., Gonzalez F., Ballester A., Blazquez J.A., Comparative study of heavy metals using different types of algae, J. Bioresour. Technol. 98 (2007) 3344–3353.

31 Saleem, M., Pirzada, T., Qadeer, R., 2007, Sorption of acid violet 17 and direct red 80 dyes on cotton fiber from aqueous solutions, J. Colloids Surf., Vol. (292), PP:246–250.

32 Sips R (1948) Combined form of Langmuir and Freundlich equations. J Chem Phys 16:490– 495

33 Stanislavsky, A. and K. Weron, K. (2013) Is there a Motivation for a Universal Behaviour in Molecular Populations Undergoing Chemical Reactions? Physical Chemistry Chemical Physics., 15, 15595–15601.

34 Sulaymon A.H., Ebrahim S.E, Abdullah S.A, Al-Musawi T.J, Removal of lead, cadmium, and mercury ions using biosorption. Desalination and Water Treatment, 24 (2010) 344–352.

35 Sulaymon A.H., Mohammed A.A., Al-Musawi T.J., Removal of lead, cadmium, copper, and arsenic ions using biosorption: equilibrium and kinetic studies. Desalination and Water Treatment 51(2013a) 4424–4434.

36 Sulaymon AH, Mohammed AA, Al-Musawi TJ. Competitive biosorption of lead, cadmium, copper, and arsenic ions using algae. Environ Sci. Pollut. Res. 20 (2013b) 3011–3023.

37 Sulaymon, A.H, Mohammed, A.A, Al-Musawi, T.J., Comparative Study of Removal of Cadmium (II) and Chromium (III) Ions from Aqueous Solution Using Low-Cost Biosorbent, Int. J. Chem. Reac. Eng., 12 (1) (2014) 1–10.

38 Yu Ji-Wei, Neretnieks, I. Single-Component and Multicomponent Adsorption equilibria on Activated Carbon of Methylcyclohexane, Toluene and Isobutyl Methyl Ketone, I& EC Research 29 (1990) 220–231.

